# The effects of exposure to predators on personality and plasticity

**DOI:** 10.1101/2020.03.26.010413

**Authors:** Amy Bucklaew, Ned Dochtermann

## Abstract

Past experiences are known to affect average behavior but effects on “animal personality”, and plasticity are less well studied. To determine whether experience with predators influences these aspects, we compared the behavior of *Gryllodes sigillatus* before and after exposure to live predators. We found that emergence from shelter and distance moved during open-field trials (activity) changed after exposure, with individuals becoming less likely to emerge from shelters but more active when deprived of shelter. We also found that plasticity in activity increased after exposure to predators and some indications that differences among individuals (i.e. “personality”) in emergence from shelter and the amount of an arena investigated increased after exposure. Our results demonstrate that experience with predators affects not only the average behavior of individuals but also how individuals differ from each other—and their own prior behavior—even when all individuals have the same experiences.

## Introduction

‘Animal personality’ has been defined as repeatable among individual variation in behavior (Dall et al. 2004). This can informally be thought of as differences amongst individuals in their average behaviors (for formal statistical definitions see Dingemanse and Dochtermann 2013). These repeatable behavioural differences have important ecological and evolutionary implications (reviewed by Wolf and Weissing 2012). Personality variation has also been found to broadly correlate with fitness (Smith and Blumstein 2008) and to be underpinned by genetic variation (Dochtermann et al. 2015).

Repeatable behavioral variation has been observed across species (Gosling 2001, Bell et al. 2009) but most research has focused on vertebrates (Bell et al. 2009), a pervasive and pernicious problem in the study of behavior (Rosenthal et al. 2017). Eusocial insects are an exception and their behavioral variation has been well described. Eusocial insects exhibit high behavioral variation due to complex caste organization, and can vary within and among colonies effect of group living on behavioral traits and hypothesizes that mixtures of behavioral types in a colony has shaped the evolution of division of labor (Jandt et al. 2014). The long-standing focus on birds and mammals (Rosenthal et al. 2017) in the study of behavior is potentially misleading as to how the majority of animals behave. For example, many invertebrates exhibit a wide range of life history events not seen in more well-studied taxa, with correspondingly unique behaviors. The dramatic physiological and morphological rearrangement accompanying metamorphosis may also lead to a rearrangement of behavior. Indeed, repeatability across even just hemimetabolous metamorphosis differs by sex in crickets (*Gryllus integer;* (Hedrick and Kortet 2012). Consequently, invertebrate personality research is necessary to determine the generality of mechanisms proposed to underpin personality variation.

As is now well recognized, the among-individual variation that characterizes personality is not mutually exclusive of the expression of behavioral plasticity. The compatibility of the two is perhaps most clear when an evolutionary reaction norm approach is adopted (Dingemanse et al. 2010) but can also be apparent with both more classic approaches to studying plasticity and by examining how variation *not* explained by among-individual differences changes with experience or time. As detailed by Westneat et al. (2015) variation unexplained by statistical models, including those used to estimate among-individual variation in studies of personality, represents plastic variation (and measurement error) and this is often a considerable amount of the biologically relevant variation in behaviors (see also Lynch and Walsh 1998, Whitman and Agrawal 2009, Berdal and Dochtermann 2019). For example, in crickets (*Acheta domesticus*) manipulation of diet during development affected plasticity/within-individual variation despite no clear effects on either average behaviors or among-individual variation in personality (Royauté and Dochtermann 2017, Royauté et al. 2019).

One particularly important potential contributor to personality and plasticity is predation risk and exposure to predators. Animals assess risks in their environment and this assessment influences decision making to optimize fitness (Lima and Dill 1990). Predation risk has been documented to shape all attributes of behavior: means, variances, and correlations. Firstly, mean behavioral responses have been shown to be affected by predator exposure. For example, in stickleback fishes an individual’s behavior while under high risk of predation is shyer and less active (Furtbauer et al. 2015). Secondly, long-term levels of variable predator exposure lead to the evolution of increased plasticity response in zooplankton, showing how behavioral variability may be influenced on an evolutionary level by predation risk (Cousyn et al. 2001). Thirdly, correlations between behaviors have been documented to change due to exposure to predators or predation risk. For example, predation strengthens behavioral correlation in stickleback as a result of both selection (bold individuals were more likely to be preyed upon) and plasticity (shift of behavior to be unaggressive; Bell and Sih 2007). Similarly, populations from environments with strong predatory pressure have been seen to have different personalities when compared to populations from safe environments (Bell 2005, Dingemanse and Reale 2005, Dingemanse et al. 2007). In contrast, Madagascar hissing cockroaches with a bold-active behavioral syndrome responded to repeated predator exposure with a shift in boldness independent of activity (McDermott et al. 2014) and a correlation between behavioral composition (proportion of bold spiders) and foraging aggressiveness in social jumping spider colonies was eliminated after exposure to predators (Wright et al. 2017).

Here we examined the effects of predator exposure on personality and plasticity in the banded cricket, *Gryllodes sigillatus*. We specifically sought to determine if mean behaviors change after exposure to predators and whether among-individual variation (i.e. personality) and within-individual variability (i.e. plasticity) are influenced by predator exposure.

## Methods

Forty adult female *Gryllodes sigillatus* were obtained from an outbred line originally collected in New Mexico (the total number of behavioral assays varied due to mortality during predator exposure). Crickets were independently housed in a 0.71-liter container with shelter, ad libitum food (commercially purchased chicken feed), and water. Crickets were housed under a 12:12h light/dark photoperiod. All testing took place during August 2018.

Individual behavior was assessed in both latency trials and open field tests conducted at the beginning of the experiment and repeated after individuals were exposed to active predators. Both latency and open field trials were conducted three times before exposure to predators and three times after. Post-exposure behavioral assays were started 24 hours after predator exposure and repeated twice at subsequent 24-hour intervals. For each behavioral test, crickets were selected at random and assigned to groups of four individuals (the testing apparatus was designed to conduct four simultaneous trials). Temperature, time, and date of each test were recorded at the start of trials.

### Latency Behavioral Assay

Crickets were randomly selected and placed into a 12 cm cylindrical shelter in the center of a 30 cm × 30cm testing arena. After a 2-minute habituation period, one end of the tube was removed and the cricket was allowed to freely emerge and move about the arena. Similar methods have been used previously to measure boldness in crickets (Hedrick 2000). Trials were recorded to accurately measure the time required for the cricket to emerge from the tube. If after 6 minutes the cricket had not emerged the trial was ended. At the end of the test, individuals were captured, their mass recorded, and were returned to housing containers. Video recordings were analyzed using VLC media player to determined amount of time until full body emergence (max 320 seconds).

### Open Field Behavioral Assay

Crickets were placed in the lower right corner of a 30cm × 30cm arena. After a 30 second habituation period, cups were removed and subjects were allowed to move freely in the testing arena for 3 minutes 40 seconds. Trials were recorded and analyzed using Ethovision (Noldus Information Technology, Wageningen, The Netherlands). Each arena was split into 32 unique zones (Figure S1). Measurements recorded were total distance moved and number of unique zones entered. At the end of the test, individuals were captured, mass was recorded, and individuals returned to their housing containers. Arenas were cleaned with 70% ethanol between tests.

### Predator Exposure

Crickets were randomly selected for the order of predator exposure. An individual cricket was trapped using a plastic cup and placed in the center of a 30 cm × 30 cm arena. All cricket handling was conducted in the same manner as for the Open Field Behavioral Assay.

Four juvenile leopard geckos (*Eublepharis macularius*) were used throughout the duration of the experiment, removed from individual housing, and placed under a cardboard cup in the same arena as were the crickets. After a 30 second habituation period, leopard geckos were released from the cups. Immediately afterwards, test subjects were released from their cups. Predator exposure experiment lasted 10 minutes or until the subject experienced an attack by the gecko. Attacks were defined as a missed strike, a strike that injured the cricket, or a successful strike. At the end of the test, individual crickets were captured, had their mass recorded, and returned to housing containers. The exposure arena was cleaned with 70% ethanol between all tests to eliminate any social or predator cues left in the arena. All geckos were housed and cared for according to NDSU IACUC Protocol A17027 and the guidelines described by the Animal Behavior Society (2012).

We did not include a control treatment because the arenas used for predator exposure were the same arenas which had previously been used for the pre-exposure open field behavioral assays. Consequently, each cricket’s pre-exposure trials act as procedural controls as the only difference pre- and post-exposure was exposure to the leopard geckos. Subsequent analyses can therefore be considered analogous to a paired t-test.

### Statistical Methods

To assess the effect of exposure to predators on behavioral means and variances we analyzed the behavioral data using linear mixed effects models (Dingemanse and Dochtermann 2013). Specifically, we fit models wherein exposure (pre or post), temperature (Celsius, mean centered), and mass (grams, mean centered) were included as fixed effects and individual was fit as a random effect.

To separately tease apart whether exposure to predators affected among- or within-individual variances (or both) we fit four distinct models with each variance differentially modeled. Following Royauté et al. (2019), we fit (i) a model where among- and within-individual variances were the same before and after exposure, (ii) a model where among-individual variances differed before and after exposure but within-individual variances remained the same, (iii) a model where within-individual variances differed but among-individual variances did not change, and (iv) a model where both among- and within-individual variances differed before and after exposure. The four models were compared using Deviance Information Criterion scores (DIC) to determine which model structure best fit the data. An alternative approach to this “character state” analysis would be using a random regression approach where the pre and post exposure periods are treated as an environmental gradient over which a reaction norm is expressed. We did not use such an approach here because for the two environment case, the character state and reaction norm approaches are mathematically equivalent. The reaction norm approach would also have made the interpretation of differences in among- and within-individual variances more difficult.

“Significance” of a fixed effect was determined based on the probability that an estimate was greater or less than zero (whichever was smaller). DIC scores were compared to each other, with particular inferential credence given to models with substantially lower scores than others. DIC scores are evaluated in a relative manner without the concrete cutoff criteria used with p-values. Instead, we interpreted a DIC difference between the best fit model and a competitor (i.e. ΔDIC) of 5 or greater as a substantial reduction of explanatory power. A ΔDIC of 10 or greater suggests a model is of little inferential utility (Barnett et al. 2010).

All models were fit using the `MCMCglmm` library (Hadfield 2010) in the R programming environment. Models were fit using parameter expanded priors and fit for 2.6 × 10^6^ iterations with a burn-in of 6 × 10^5^ iterations and a 2000 iteration thinning interval. This combination of priors and chain length maintained high levels of mixing and low levels of autocorrelation (variance component autocorrelations were all less than 0.06 even for Poisson distributed behaviors). Distance moved was fit as Gaussian distributed while unique zones visited and emergence (no or yes) were fit, respectively, as Poisson and Binary distributed. From these models we also estimated the behavioral variance explained due to fixed effects and estimated adjusted repeatabilities following Nakagawa and Schielzeth (2010) & (2013).

We were not able to fit multivariate models due to the mixture of error distributions: multivariate models failed to properly converge despite doing so for univariate models. This prevented us from assessing the effects of exposure to predators on behavioral syndromes despite that originally being one question of interest.

## Results

Prior to exposure, mean distance traveled during the open field trials was 220.43 cm, mean number of unique zones visited 15.51, and mean latency to emerge was 164.10 seconds. Post-exposure mean distance moved was 305.49 cm, mean number of unique zones visited 15.95, and mean latency to emerge 245.02 seconds. Activity in the open field and emergence from shelter significantly differed pre to post exposure (p = 0.034 & 0.006 respectively; Figure 1, Table 1). Unique Zones visited were not significantly different pre to post exposure (Figure 1B, Table 1). Mass significantly affected activity (p = 0.03) and UZ (p = 0.03), but not emergence (p = 0.956). Temperature did not have a significant interaction with exposure conditions on any behavioral responses (Table 1).

**Table 1.**
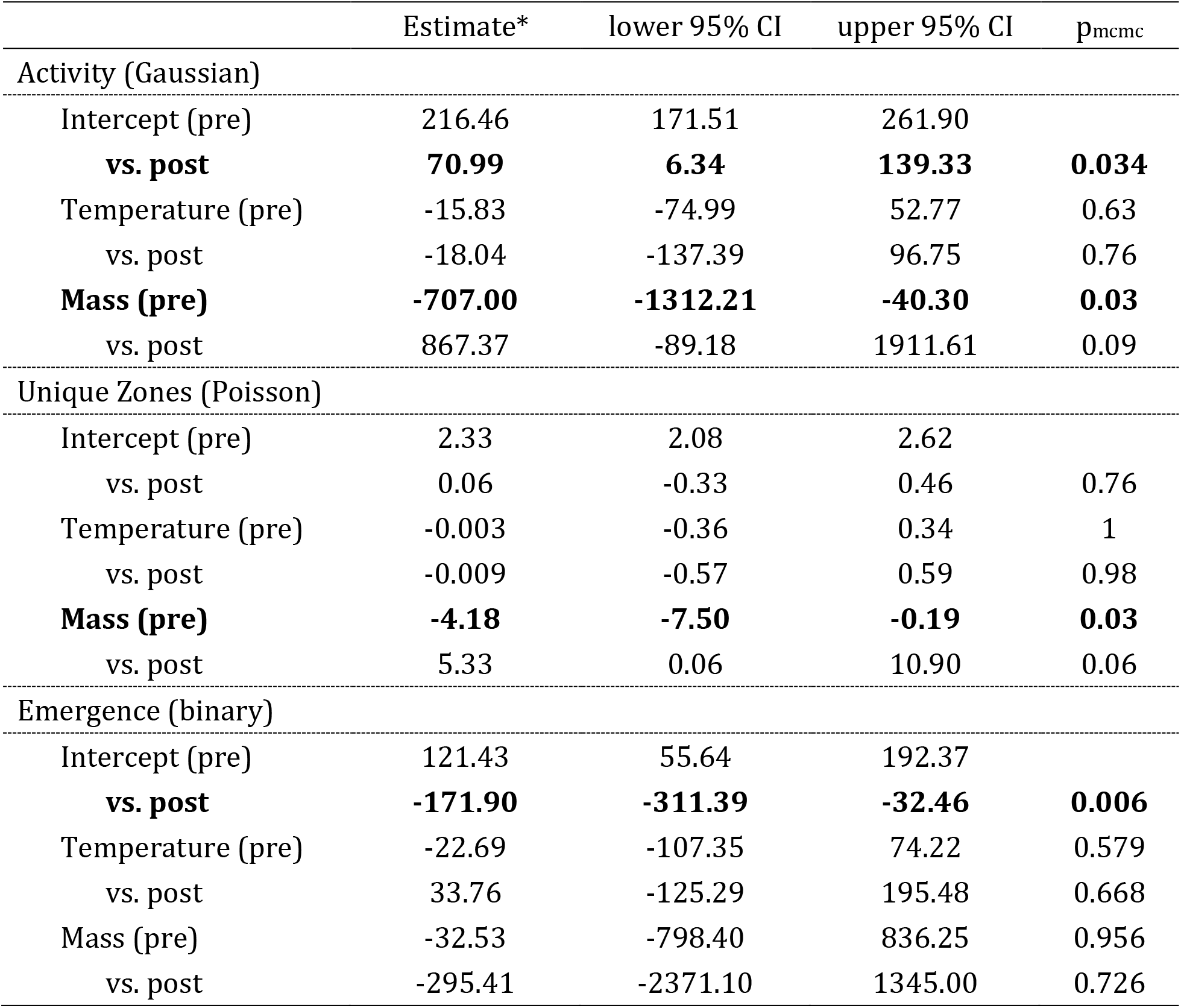
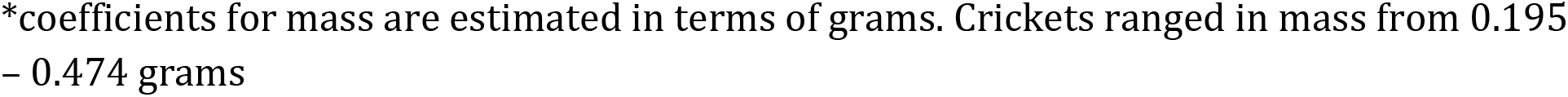
Summary of fixed effect estimates on behavioral responses, values are posterior means. Results are from Bayesian analyses fit using the `MCMCglmm` package in R. Fixed effect structure was the same for each of the three behavioral responses: Pre-exposure was fit as the intercept against which the estimate for post-exposure was contrasted (i.e. the estimate for Post-exposure would be the sum of the pre- and post-exposure estimates). The effects temperature (Celsius, mean centered) and mass (g, mean centered) were also estimated for post-exposure, with pre-exposure then contrasted (i.e. exposure × temperature and exposure × mass interactions). Activity was fit using a linear mixed-effects model while the unique zones visited and whether an individual emerged from shelter were fit with generalized linear mixed-effects models as Poisson distributed and as a categorical (binary) outcome respectively. Random effects structure was as in the best fitting model (Table 2). P_mcmc_ for a factor is estimated based on the number of posterior estimates that did not exclude 0.

| | Estimate* | lower 95% CI | upper 95% CI | $p_{\text{mcmc}}$ |
| --- | --- | --- | --- | --- |
| Activity (Gaussian) |  |  |  |  |
| Intercept (pre) | 216.46 | 171.51 | 261.90 |  |
| **vs. post** | **70.99** | **6.34** | **139.33** | **0.034** |
| Temperature (pre) | -15.83 | -74.99 | 52.77 | 0.63 |
| vs. post | -18.04 | -137.39 | 96.75 | 0.76 |
| **Mass (pre)** | **-707.00** | **-1312.21** | **-40.30** | **0.03** |
| vs. post | 867.37 | -89.18 | 1911.61 | 0.09 |
| Unique Zones (Poisson) |  |  |  |  |
| Intercept (pre) | 2.33 | 2.08 | 2.62 |  |
| vs. post | 0.06 | -0.33 | 0.46 | 0.76 |
| Temperature (pre) | -0.003 | -0.36 | 0.34 | 1 |
| vs. post | -0.009 | -0.57 | 0.59 | 0.98 |
| **Mass (pre)** | **-4.18** | **-7.50** | **-0.19** | **0.03** |
| vs. post | 5.33 | 0.06 | 10.90 | 0.06 |
| Emergence (binary) |  |  |  |  |
| Intercept (pre) | 121.43 | 55.64 | 192.37 |  |
| **vs. post** | **-171.90** | **-311.39** | **-32.46** | **0.006** |
| Temperature (pre) | -22.69 | -107.35 | 74.22 | 0.579 |
| vs. post | 33.76 | -125.29 | 195.48 | 0.668 |
| Mass (pre) | -32.53 | -798.40 | 836.25 | 0.956 |
| vs. post | -295.41 | -2371.10 | 1345.00 | 0.726 |

\*coefficients for mass are estimated in terms of grams. Crickets ranged in mass from 0.195 – 0.474 grams

**Figure 1:**
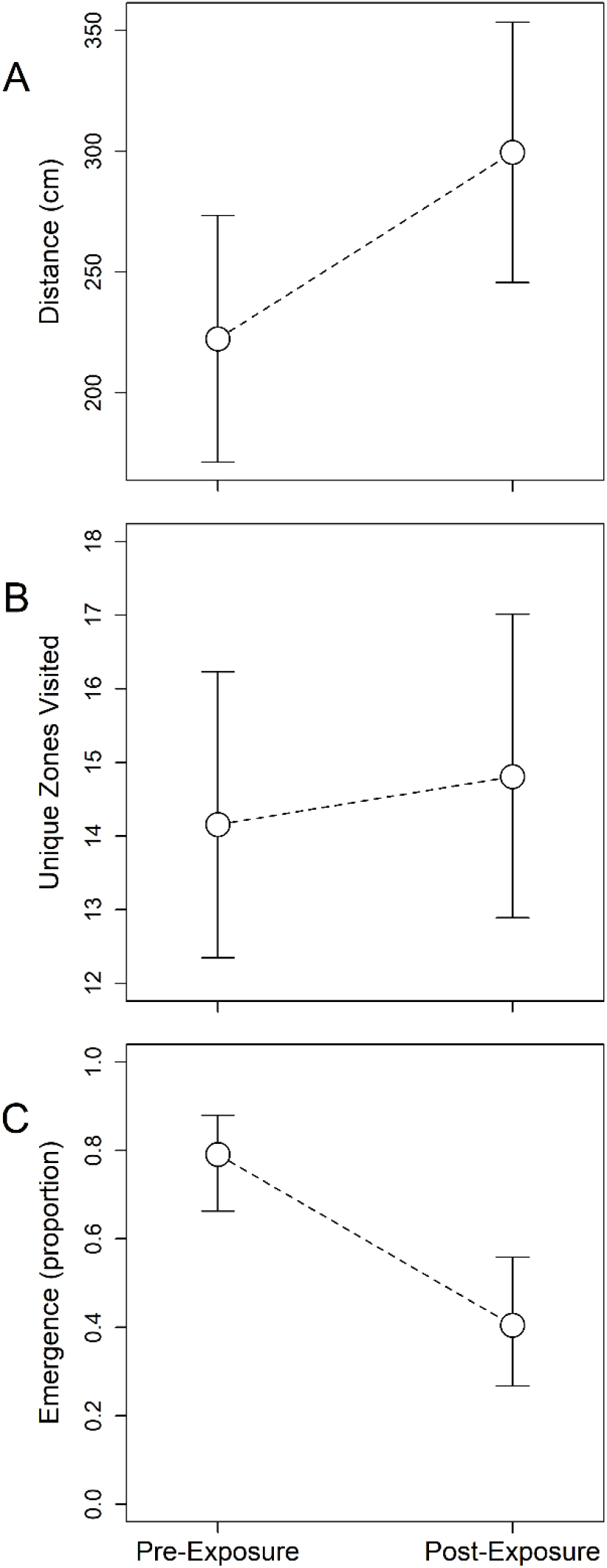
Differences in average behavioral response pre- and post-exposure to a predator. Points are the posterior modal estimates and error bars show the 95% highest probability density intervals. (A) Activity, estimated as distance traveled in the open field, significantly increased following exposure (Pmcmc = 0.034). (B) The number of unique zones visited did not significantly change in response to predator exposure. (C) Probability of emergence significantly decreased following exposure (Pmcmc = 0.006).

The influence of predator exposure on among-individual and within-individual variances of behavioral response varied by behavior (Table 2). For activity, the top model suggested that within-individual variation, but not among-individual variation, differed following exposure to predators (Table 2). For UZ, the top model included differences in among-individual variances but no one model could be distinguished from the rest (Table 2). Similarly, for emergence, the top model included a change in among-individual variance following exposure but a similarly weighted model suggested no change in variances (Table 2).

These model comparison conclusions about differences in variances pre- and post-exposure are reflected in changes in both the magnitude of variances and in the magnitudes of repeatabilities (*τ*; Table 3). Most notably, the repeatability of emergence drastically increased following exposure—from 0.13 to 0.59. This was driven by an increase in the among-individual variance following exposure (Table 3), a finding consistent with the top model examining changes in variances. Recall, however, that this top model could not be clearly distinguished from others (Table 2). In contrast, based on model comparison results and estimate uncertainties, an increase in repeatability of activity following exposure was driven by a substantial decrease in within-individual variances.

**Table 2.** Results from model comparisons evaluating the fit of models in which either, or neither, among-individual (*V*_*i*_) and within-individual variances (*V*_*w*_) of behavioral responses differ before and after exposure to predators. Models were compared based on Deviance Information Criteria scores (DIC) and the difference in DIC between the best fitting model, i.e. the model with the lowest DIC score (in bold), and each other model (ΔDIC). Models within 2 ΔDIC values do not substantively differ in fit and are indicated by italicization

| Model | Variance comparison | Activity |  | Unique zones |  | Emergence |  |
| --- | --- | --- | --- | --- | --- | --- | --- |
| | | DIC | ΔDIC | DIC | ΔDIC | DIC | ΔDIC |
| Model 1 | $V_i = \& V_w =$ | 3087.76 | 8.98 | *1397.15* | *0.26* | *5.46* | *0.32* |
| Model 2 | $V_i \neq \& V_w =$ | 3085.42 | 6.64 | **1396.89** | **0** | **5.14** | **0** |
| Model 3 | $V_i = \& V_w \neq$ | **3078.78** | **0** | *1397.19* | *0.3* | 14.52 | 9.38 |
| Model 4 | $V_i \neq \& V_w \neq$ | 3084.37 | 5.59 | *1397.11* | *0.22* | 7.39 | 2.25 |

**Table 3.** Among-individual variances (*V*_*i*_) and within-individual variances (*V*_*w*_) for each of the three behavioral responses, estimated separately pre- and post-exposure. Also reported is the amount of variation explained by fixed effects (*V*_*F*_) and the unadjusted repeatability (*τ)*.

| | $V_i$ (CI) | $V_w + \text{DSV}^*$ (CI) | $V_F$ (CI) | $\tau^{**}$ (CI) |
| --- | --- | --- | --- | --- |
| Activity |  |  |  |  |
| Pre-exposure | 4259<br>[1 : 12932] | 25770<br>[20618 : 37871] | 3334<br>[699 : 7971] | 0.14 [0 : 0.31] |
| Post-exposure | 15155<br>[0 : 33054] | 47689<br>[35750 : 68101] |  | 0.20 [0 : 0.41] |
| Unique zones |  |  |  |  |
| Pre-exposure | 0.28 [0 : 0.58] | 0.60 [0.48 : 1.04] | 0.05<br>[0.01 : 0.12] | 0.24 [0 : 0.47] |
| Post-exposure | 0.22 [0 : 0.61] | 0.92 [0.68 : 1.50] |  | 0 [0 : 0.38] |
| Emergence |  |  |  |  |
| Pre-exposure | 128<br>[0 : 20079] | 16744<br>[678 : 49224] | 3056<br>[494 : 19387] | 0 [0 : 0.42] |
| Post-exposure | 14879<br>[638 : 104923] | 1913<br>[54 : 39224] |  | 0.65 [0.33 : 0.86] |

\*DSV: Distribution specific variance. Calculated following Nakagawa and Schielzeth (2010): for Poisson distributed data (Unique zones), calculated as $\ln\left(\frac{1}{e^{\beta_0}} + 1\right)$ ; for binary data, calculated as: $\frac{\pi^2}{3}$
\*\* $\tau$ : unadjusted repeatability. Calculated following Nakagawa and Schielzeth (2010) as: $\frac{V_i}{V_i + V_F + V_w}$ ; includes DSV.

## Discussion

Comparing pre- and post-predator exposure behavior, we found a shift in both mean behaviors and individual variances. Notably, after predator exposure, activity significantly increased and emergence significantly decreased in *Gryllodes sigillatus* (Table 1, Figure 1A). Biologically, increased activity can be used as an anti-predator response (Jones and Godin 2009, Wilson et al. 2010) and crickets are known to respond by increasing activity in response to some spider predators (Binz et al. 2014). Similarly, in grasshoppers, individuals respond to predators with active hunting strategy (as is the case for leopard geckos) by increasing their activity (Miller et al. 2014). In this study, subjects potentially increased activity after predator exposure as a general response to avoid capture by predators. Similarly, delaying emergence (Figure 1C) is often considered a measure of cautiousness, or the inverse of boldness (Hedrick and Kortet 2012). Consequently, the decreased probability of emergence following exposure is perhaps an anti-predator response in which crickets avoid leaving safety when there exists a known predator threat.

Interestingly, exposure to predators did not affect behavioral variation in the same direction across behaviors. Plasticity, or within-individual variation, differed pre- and post-exposure for the distance moved (Table 2) and was greater after exposure (Table 3). Therefore, individuals become more plastic in their behavior after exposure to predators. This also creates the possibility for increased individual differences in behavioral responses to predator interaction with some individuals moving erratically and others not at all as an anti-predator tactic. Predator-exposure coping mechanisms are therefore potentially individual specific and successful tactics can lead to evolutionary changes. Unfortunately, assessing this possibility is not possible with the current data, requiring the use of data-hungry approaches (such methods are described in Cleasby et al. 2015).

Among-individual variance also potentially differed pre- and post-exposure for emergence (Table 2), being greater after exposure (Table 3). While there was mixed support for this conclusion, it suggests that individuals may become more fixed in their differences from each other in whether they emerge or not after exposure. Further, repeatability of the behavior increased substantially despite an absence of clear specific differences in among- or within-individual variances. These results suggest an increased conformity to an individual’s behavioral type after exposure, i.e. anti-predator tactics are individual specific. Importantly this also shows that the behavior of individuals continues to differ—and possibly increases—even when presented with the same experiences. This could suggest the emergence of new behavioral tendencies: if individuals randomly emerge from shelter under conditions without pressure, new individually specific latency personalities may effectively canalize after life experiences. This possibility is consistent with Bayesian updating models of development (Stamps et al. 2018) but, due to the ambiguous results regarding emergence here, future work should further investigate this possibility in crickets.

Previous research has similarly shown that predator exposure changes average anti-predator responses from subjects, with putative effects on survival probabilities. For example, predator-naïve mammals exposed to predators under controlled conditions showed changes in behavior including increased wariness and greater flight initiation distances (West et al. 2018). Second-hand fear cues, bedding material used by predator-exposed voles, lead to increased litter sizes and sex based behavioral changes in regard to the bedding (Haapakoski et al. 2018). Such results show a population level effect on individuals who have not been directly exposure to a predator. In contrast, the results of our study suggest that patterns of behavioral change rely on individual specific anti-predator responses and a combination of changes in average behavior and in variances. This result has important implications for our understanding of the interaction between selection and personality. For example, in crabs (*Panopeus berbstii*), individual specific behavioral responses interacted with predator type to affect survival (Belgrad and Griffen 2016).

Animals constantly assess risks in their environment and use this information to guide decision making (Lima and Dill 1990), with personality having been shown to affect survival rates in response to predation threat (Santos et al. 2015). The results of this study show that *Gryllodes sigillatus* likewise use information from predator experience to shape behavior. Comparing pre- and post-exposure behavioral assays, we demonstrated a shift in mean behaviors and in how individuals differ from each other (variances). Thus, exposure to predators can change among-individual differences—personality—and plasticity.

## Acknowledgements

The authors thank Raphaël Royauté, Monica Berdal, Jeremy Dalos, Jenna LaCoursiere, Ishan Joshi, and Brady Klock for with cricket rearing, advice, and comments. We also thank Matt Smith for assistance with the leopard geckos and the National Science Foundation and the National Science Foundation’s Research Experience for Undergraduates program for funding (NSF-IOS 1557951).

---

**Figure S1:**
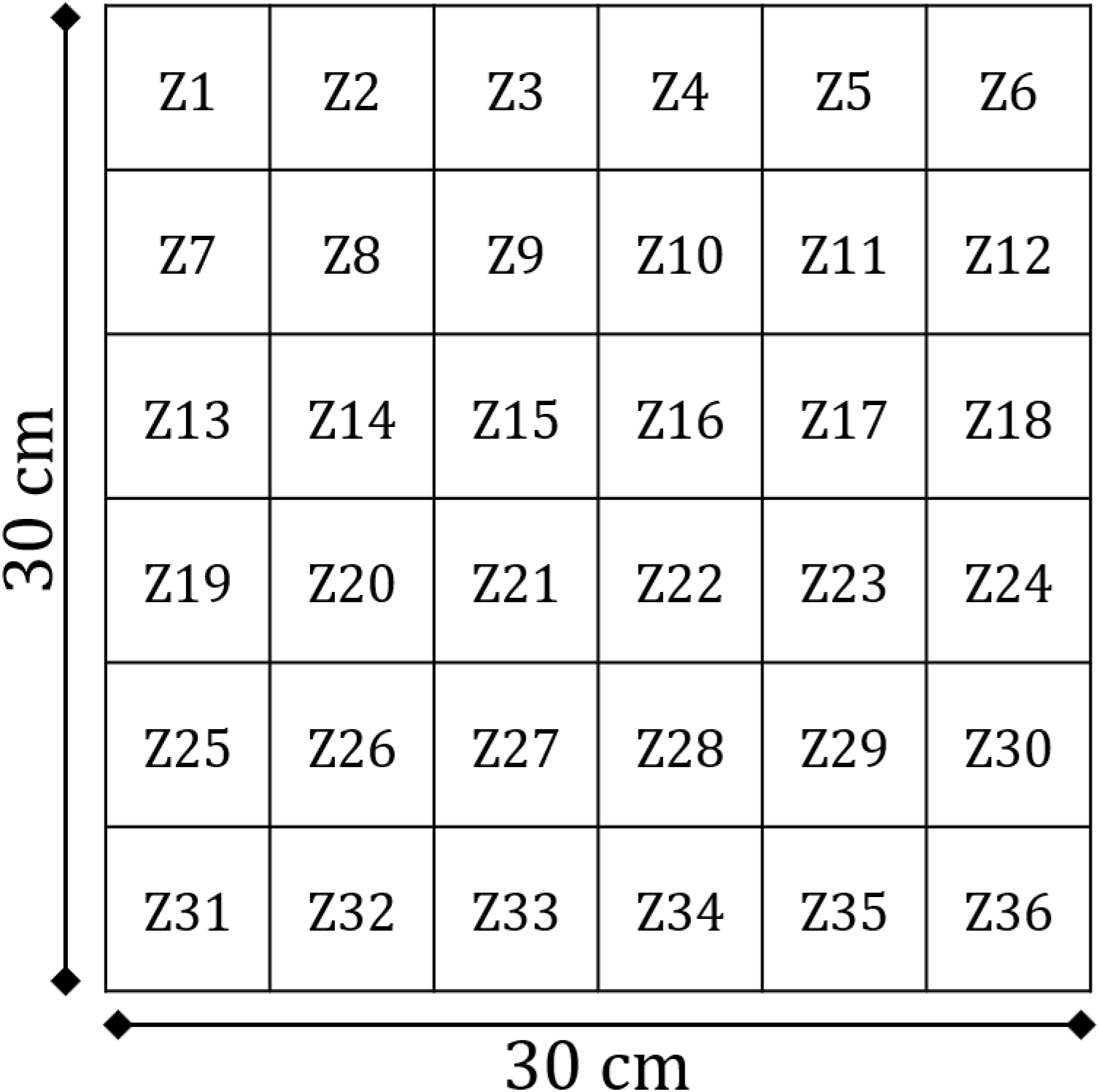
Schematic of the open-field arena.

